## Supplement_Quant_Paper.pdf for "Precursor intensity-based label-free quantification software tools for proteomic and multiomic analysis within the Galaxy Platform"

### Supplementary Figures:

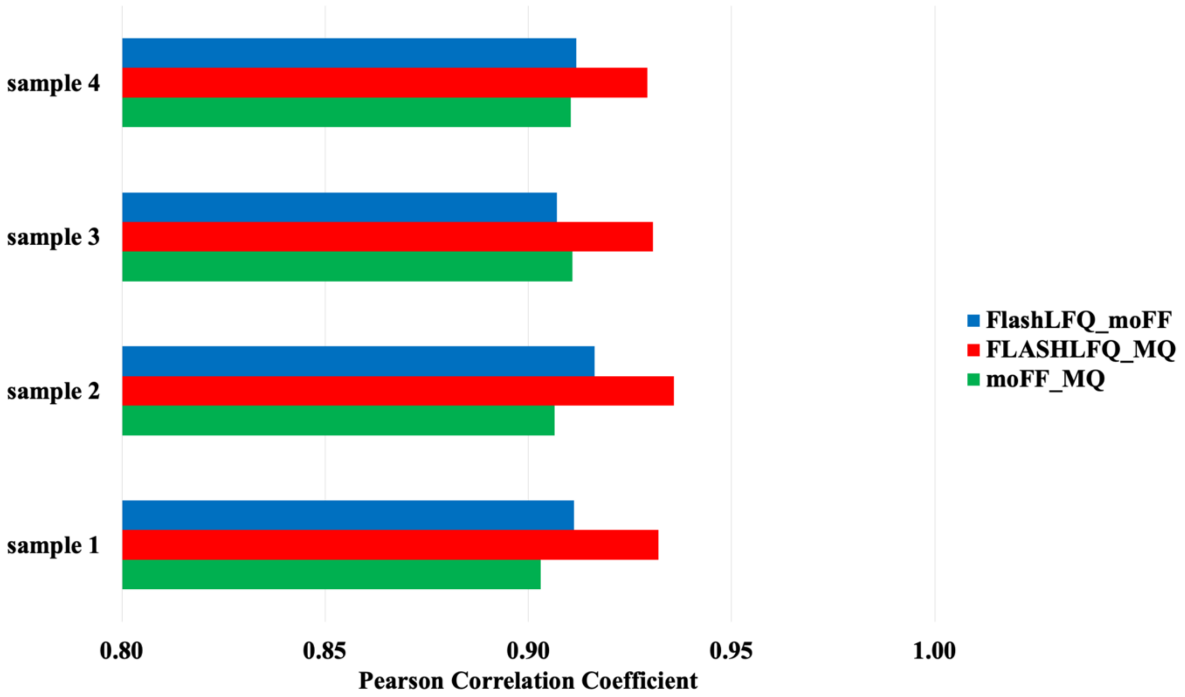


**Supplementary Figure S1: Peptide Correlation :** The raw intensities of the ABRF peptides were correlated using Pearson Correlation coefficient. Output from FlashLFQ and moFF correlated well with MaxQuant.

**(a)**

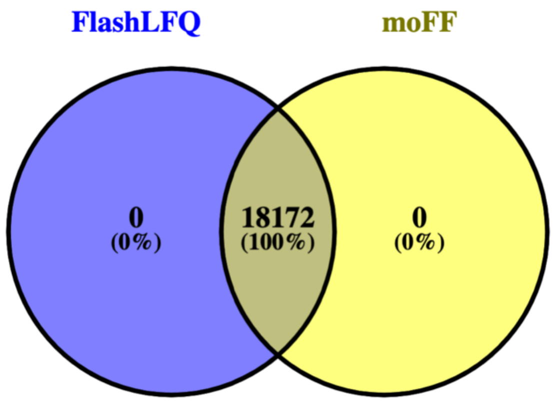


**(b)**

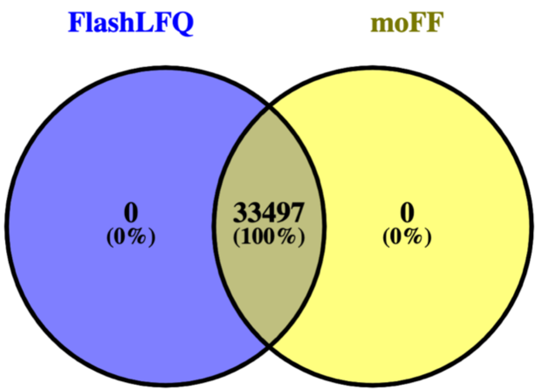


**Supplementary Figure S2: Peptide Overlap across FlashLFQ and moFF :** A Venn diagram of the quantified ABRF (a) and UPS (b) unique peptides are shown to display the coverage of the peptides across FlashLFQ and moFF. The input for both the tools was the PSM report from the Peptide shaker containing 18172(ABRF) and 33497 (UPS) unique peptides.

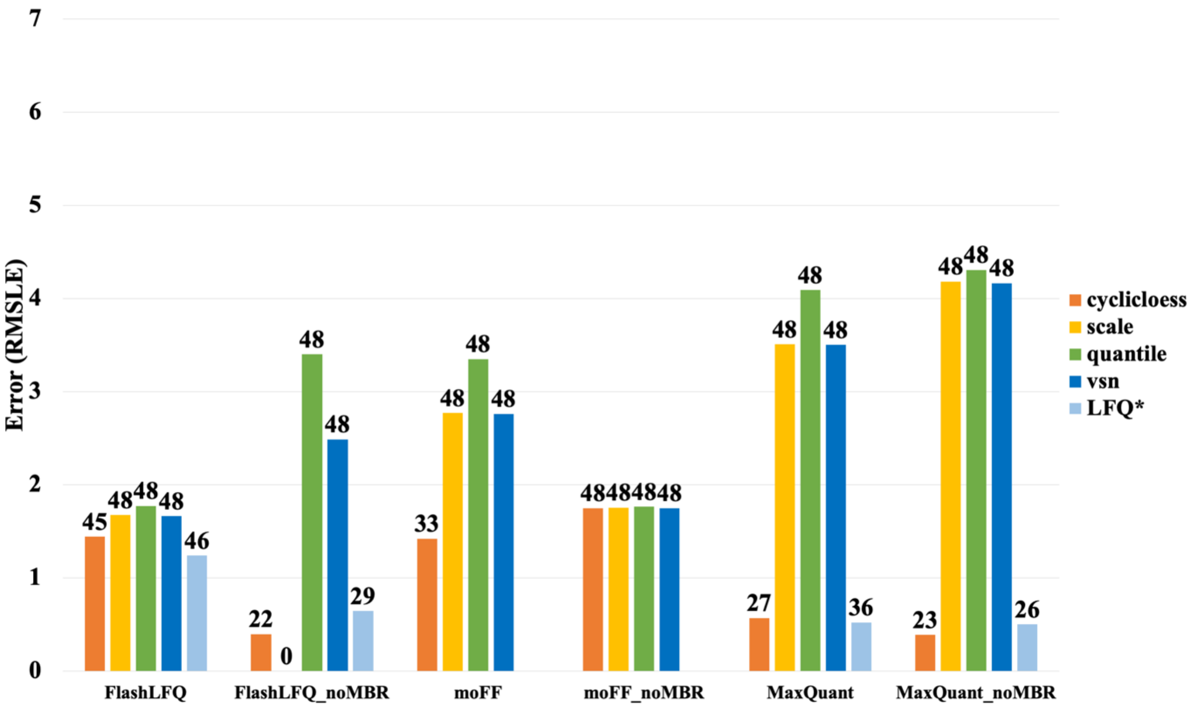

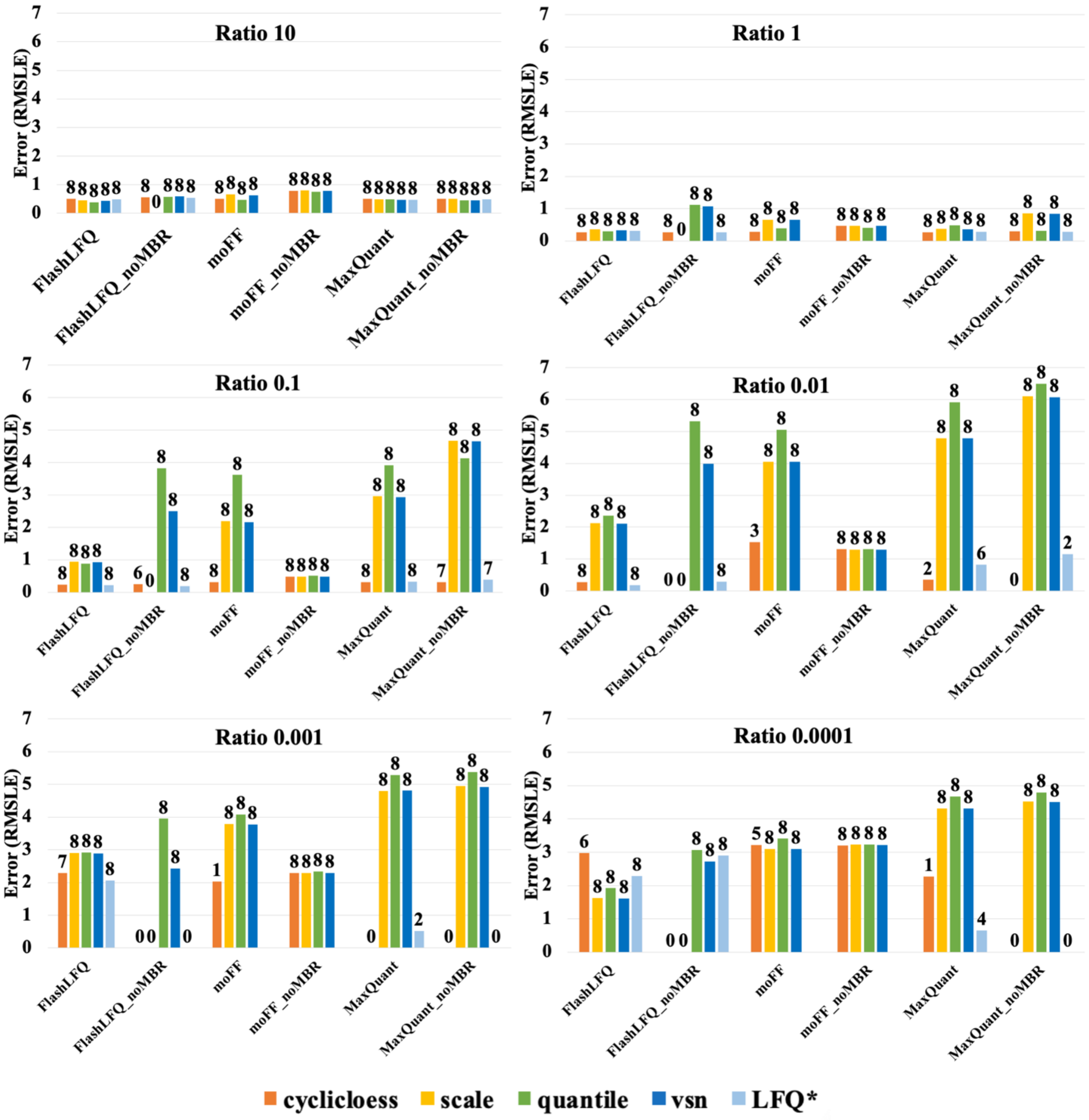


**Supplementary Figure S3A) Fold-change accuracy (MBR vs no-MBR) of all proteins:** After normalization, the estimated protein ratios for all the identified UPS proteins were compared to the true ratios, using the Root mean squared log error (RMSLE). The plot represents the comparison between MBR and no-MBR using normalization methods. Note that LFQ values represent MaxQuant's and FlashLFQ's normalized value. The value on the top of the bars denotes the number of proteins that were quantified. Although the bar graph shows that MaxQuant's MaxLFQ performs the best compared to all, we notice that the number of UPS proteins identified and used for the calculation were less compared to other tools.

**Supplementary Figure S3B) Fold change accuracy (MBR vs no-MBR) of proteins with similar estimated ratios:** In total there are 48 UPS proteins, we classified the UPS proteins into different groups based on the UPS2/UPS1 ratio estimation, the true ratios run from 10 to $10^{-4}$ . Value on the top of the bars denote the number of proteins that were quantified using each normalization method. This is a comparative study between MBR and no MBR. The RMSLE of the intensity ratio was used to measure the accuracy of the estimated fold change. The figure shows that moFF and FlashLFQ work well in comparison to MaxQuant for ratio estimation of low abundance proteins(<0.1).

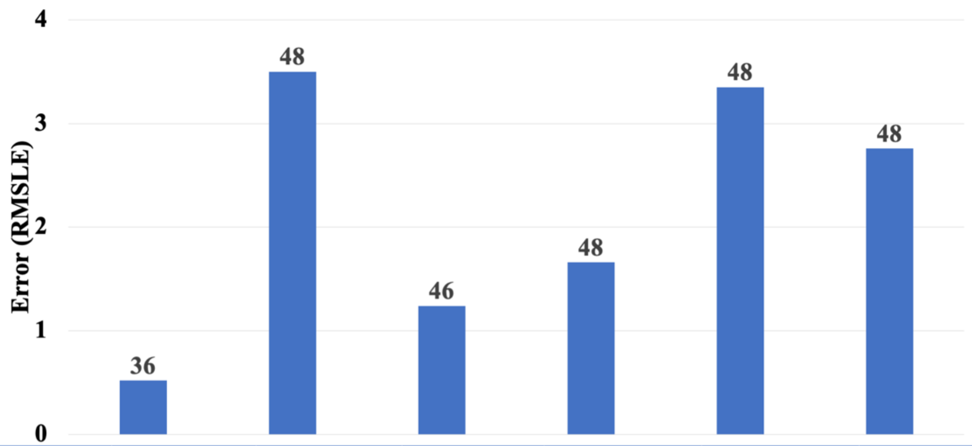


| Search Engine | MaxQuant | MaxQuant | SG/PS* | SG/PS* | SG/PS* | SG/PS* |
|---|---|---|---|---|---|---|
| **Peptide Quantification** | MaxQuant | MaxQuant | FlashLFQ | FlashLFQ | moFF | moFF |
| **Normalization** | MaxQuant | vsn | FlashLFQ | vsn | quantile | vsn |
| **Protein Quantification** | MaxQuant | PECA | FlashLFQ | PECA | PECA | PECA |

**Supplementary Figure S4) Fold Accuracy of Tools with inbuilt and external normalization and protein quantitation:** This figure displays the errors in Fold change accuracy while using inbuilt and external normalization and protein level quantitation features of the evaluated tools.
